# Obesity Facilitates Sex-Specific Improvement In Cognition And Neuronal Function In A Rat Model Of Alzheimer’s Disease

**DOI:** 10.1101/2024.01.11.575200

**Authors:** Aaron Y. Lai, Dustin Loren V. Almanza, Jessica A. Ribeiro, Mary E. Hill, Matthew Mandrozos, Margaret M. Koletar, Bojana Stefanovic, JoAnne McLaurin

## Abstract

Obesity reduces or increases the risk of developing Alzheimer’s disease (AD) depending on whether it is assessed in mid-life or late-life. There is currently no consensus on the relationship between obesity and AD or the mechanism or their interaction. Here, we aim to differentiate the cause-and-effect relationship between obesity and AD in a controlled rat model of AD. We induced obesity in 9-month-old TgF344-AD rats, that is pathology-load wise similar to early symptomatic phase of human AD. To more accurately model human obesity, we fed both TgF344-AD and non-transgenic littermates a varied high-carbohydrate-high-fat diet consisting of human food for 3 months. Obesity increased overall glucose metabolism and slowed cognitive decline in TgF344-AD rats, specifically executive function, without affecting non-transgenic rats. Pathological analyses of prefrontal cortex and hippocampus showed that obesity in TgF344-AD rats produced varied effects, with increased density of myelin and oligodendrocytes, lowered density and activation of microglia that we propose contributes to the cognitive improvement. However, obesity also decreased neuronal density, and promoted deposition of amyloid-beta plaques and tau inclusions. After 6 months on the high-carbohydrate-high-fat diet, detrimental effects on density of neurons, amyloid-beta plaques, and tau inclusions persisted while the beneficial effects on myelin, microglia, and cognitive functions remained albeit with a lower effect size. By examining the effect of sex, we found that both beneficial and detrimental effects of obesity were stronger in female TgF344-AD rats indicating that obesity during early symptomatic phase of AD is protective in females.

## INTRODUCTION

Worldwide total dementia cases are expected to rise to more than 150 million by 2050 (Nichols et al., 2022). Alzheimer’s disease (AD) is the most common form of dementia and is currently without preventative or disease modifying treatments (Association, 2019). Identifying modifiable risk factors is therefore an area of significant importance. Obesity has garnered much attention as a correlate of dementia risk, yet its precise relationship with AD has been a highly contested topic: Obesity during mid-life or the prodromal phase of AD is associated with increased risk of developing or early onset of the disease (Kivipelto et al., 2005; Whitmer et al., 2007; Beydoun et al., 2008; Hassing et al., 2009; Tolppanen et al., 2014; Pedditizi et al., 2016); however late-life obesity is paradoxically associated with lower incidence of AD (Luchsinger et al., 2007; Atti et al., 2008; Fitzpatrick et al., 2009; Tolppanen et al., 2014; Pedditizi et al., 2016; Sun et al., 2020). Two explanations have been proposed for this apparent ‘obesity paradox’: 1) Weight loss that occurs during the prodromal phase of AD is a consequence rather than a cause such that weight maintenance and obesity is a result of intact cognitive health. 2) Body composition of obese elderly individuals offers protection against cognitive decline.

To differentiate the cause-and-effect relationship between obesity and cognitive maintenance, studies have experimentally induced obesity using high-fat diets in controlled transgenic mouse models of AD. These studies found that obesity could be both detrimental and protective to cognitive maintenance in the presence of AD transgenes (reviewed in Amelianchik et al., 2022), thus adding further intrigue to the ‘obesity paradox’. Here, we propose a novel approach that overcomes several key limitations of previous obesity models. We induced obesity in the TgF344-AD (TgAD) rat model by feeding them a high-carbohydrate, high-fat diet (HCHF) consisting of varied human food that has been shown to continuously engage the rats in the diet (Gomez-Smith et al., 2016). This mimics human obesity more accurately than gaining weight on a non-varied diet. Secondly, the TgF344-AD rat model develops neuronal loss in addition to cognitive deficits, parenchymal and vascular Aβ deposition, and tau inclusions (Cohen et al., 2013; Morrone et al., 2020, 2022) which mirrors clinical AD more closely than do mouse models of AD. Furthermore, we induced obesity at an age with mild cognitive and pathological impairment resembling early symptomatic stages of AD (Morrone et al., 2022). This particular timing allows for clearer differentiation that the observed effects are not due to underlying mechanisms during the prodromal phase. As past clinical association studies have found that the relationship between body weight and cognitive impairment is sex-specific (Joo et al., 2018; Noh et al., 2019), we also examined sex differences which have been a heavily overlooked aspect in animal studies of AD.

In the current study, we aim to determine if obesity developed during early stage of AD is protective against cognitive decline, which cellular mechanisms contribute to this protection, and whether the effect is sex-specific. We induced obesity in the TgAD rats at 9 months of age where they develop cognitive and pathological deficits similar to early symptomatic stages AD (Morrone et al., 2022). Both non-transgenic (NTg) and TgAD rats were fed with HCHF to increase their body mass index (BMI). After either 3- or 6-months on HCHF, we assessed their brain metabolism, cognitive performance, progression of AD pathology, and overall health of neurons and glial cells. Longitudinal HCHF treatment allows us to determine effects of BMI across AD progression and disaggregation of males and females allow detection of sex-specific differences.

## MATERIALS AND METHODS

### Animals

Non-transgenic F344 rats (NTg) and TgF344-AD rats (TgAD) harboring APP-Swedish and PSEN1-deltaE9 mutations (Cohen et al., 2013) were housed on a 12 hour:12 hour light/dark cycle with each cage housing two cage-mates. All cages receive food and water *ad libitum* and daily monitoring for health and normal grooming behaviors. The Animal Care Committee of the Sunnybrook Health Sciences Center, which adheres to the Policies and Guidelines of the Canadian Council on Animal Care, the Animals for Research Act of the Provincial Statute of Ontario, and the Federal Health of Animals Act has granted ethical approval of all animal experiments outlined in this study.

At 9 months of age, NTg and TgAD rats of both sexes were randomly assigned to one of two diet groups: Standard chow (CHOW) or diet enhanced with high carbohydrates and high fat (HCHF). To induce obesity, HCHF rats were given the choice between standard chow and three highly palatable human HCHF items *ad libitum* as outlined previously (Gomez-Smith et al., 2016). The HCHF rats were also given the choice between water and 12% sucrose solution. The three HCHF items were rotated with three different HCHF items three times a week to maintain appetite and palatability. Weight gain for each rat is monitored weekly until sacrifice at 12 and 15 months of age. BMI was expressed as weight/length (grams/cm2) as described previously (Novelli et al., 2007; Rabiu et al., 2017) where length was measured from nose to anus. Statistical significance was measured using one-way ANOVA followed by Tukey’s *post-hoc*.

### Magnetic Resonance Spectroscopy (MRS)

MRS data were obtained using Bruker BioSpec 7T before and after 2-deoxy-glucose injection to assess alterations in glucose and other metabolites as described previously (Joo et al., 2022). Metabolic profile of a single voxel (4 x 2.0 x 1.5 mm3) in the left hippocampus (HIPP) was assessed using single voxel point resolved spectroscopy (PRESS) sequence (TR/TE = 2500/16 ms, 2048 spectral data points, bandwidth of 4 kHz and with 1000 averages). The voxel placement (centered at −4.0 mm AP and 3.0 mm ML) was adjusted to attain water peak linewidth below 12 Hz after localized shimming. Variable pulse power and optimized relaxation delays approach was used for water suppression. Brain metabolite concentrations were quantified in the frequency domain using a basis set for a short TE PRESS sequence (Landheer et al., 2021) in LCModel (Provencher, 2001). For exclusion criteria of metabolite concentrations, a cutoff of 30% absolute Cramer–Rao lower bounds (CRLB) averaged across all scans was used where individual samples with high CRLB are included to not bias the mean estimated concentration of cohort data (Kreis, 2016; Fowler et al., 2021). For each metabolite, the means and standard deviation of absolute CRLB were used to quality filter the estimated metabolite concentrations. Estimated metabolite concentration with absolute CRLB within the mean ± 2 x standard deviation were considered in the subsequent analysis (Kreis, 2016). Statistical significance was measured using linear mixed effects in R.

### Barnes Maze

Cognitive functions were assessed at 12 and 15 months of age after 3 and 6 months of dietary treatment using Barnes Maze as described previously (Lai et al., 2022; Morrone et al., 2022). In brief, after the rats completed the learning across three days with two trials per day. One probe trial was conducted three days later to assess spatial memory. The following day, reversal trials began for five days (two trials per day) in which the escape hole location was altered to measure executive function. EthoVision was used to record videos of each rat during the trials which were analyzed by a blind investigator. Statistical significance was measured by either two-way (probe trial) or three-way (learning and reversal trials) ANOVA computed using GraphPad Prism.

### Immunofluorescence and immunohistochemistry

At end of treatment after behavior and MRS experiments, rats 12 and 15 months of age were transcardially perfused under 5% isoflurane with PBS-0.1% heparin followed by PBS-4% paraformaldehyde, overnight post-fixation in PBS-4% paraformaldehyde, then cryopreservation in PBS-30% sucrose for microtome sectioning. Three evenly spaced 40 µm sections were sampled between +3.5 and +2.0 mm AP (prefrontal cortex, PFC) as well as between −3.0 and −4.5 mm AP (HIPP).

For immunofluorescence, floating sections were blocked in PBS containing 0.5% Triton X-100 and 0.5% bovine serum albumin for 1 hour, then incubated overnight with the following primary antibodies diluted in block: Anti-myelin basic protein (MBP, 1:100, Millipore #MAB386), anti-NeuN (1:500, Millipore #ABN90), anti-Iba1 (1:500, Wako #019-19741) followed by 2 hour probing with the appropriate secondary antibodies diluted in block: Anti-guinea pig IgG-Alexa 594 (1:250, ThermoFisher #A11076), anti-rat IgG-Alexa 488 (1:250, ThermoFisher #A21208), Anti-rabbit IgG-Alexa 488 (1:250, ThermoFisher #A21206), before mounting.

For immnohistochemistry of PDGFR-α (1:50, R&D Systems #AF1062) and Olig2 (1:250, Abcam #ab109186), floating sections were first quenched in 3% H_2_O_2_-PBS for 30 minutes then subjected to 45 minutes of heating in 85°C in pH 6.0 10 mM sodium citrate before the same blocking, primary, and secondary antibody (1:500 anti-goat IgG-biotin, JacksonImmuno #705-065-147; 1:400 anti-rabbit IgG-biotin, Vector #PK-4001) labeling as outlined above, followed by one hour incubation with ABC kit (1:400 A and 1:400 B, Vector #PK-4001), before dehydration and mounting.

Immunohistochemistry for Aβ plaques and tau inclusions were performed as previously described (Morrone et al., 2022).

Myelin staining was carried out using Black Gold II (HistoChem #2BGII). Saline-washed sections were mounted, then incubated with 1 mL of 0.3% Black Gold II-saline on a heating block at 60°C for 6 minutes, washed in distilled water then 1% sodium thiosulfate-saline. The slides were washed in distilled water before dehydration and mounting.

Zeiss Observer.Z1. was used to acquire labeled images which were quantitatively analyzed using ImageJ. The investigator is blinded to treatment conditions throughout the analysis. Threshold for each analyzed image was set to where the background contains 7-10 pixels per 150 µm^2^. For measuring density of myelin, NeuN, Iba1, MBP, and Aβ plaques, particles were analyzed without size exclusions. For automated cell counting of PDGFR-α and Olig2, only particles 20-200 pixels in size were included. For analysis of microglial morphology, 40 microglia were randomly sampled and classified into either ramified (0), hypertrophic (1), bushy (2), or amoeboid (3) according to previously described criteria (Wyatt-Johnson et al., 2017). Overall microglial activation is represented by the sum of each cell type multiplied by the bracketed value. Counting of PHF-1-positive inclusions was carried out manually. Statistical significance was measured by two-way ANOVA followed by Tukey’s *post-hoc* or t-test where there are no multiple comparisons and computed using GraphPad Prism.

## RESULTS

### HCHF markedly increased BMI of both NTg and TgAD rats

We first set out to determine the extent to which HCHF regiment induced obesity in our rats. We fed NTg and TgAD rats for 3- and 6-months with either normal CHOW or HCHF diet starting at 9 months of age. Although an imperfect measure, BMI is commonly utilized as a measure of obesity in humans and thus was utilized in our study.

Male NTg and TgAD rats fed normal CHOW diet had similar starting BMIs (p=0.121 N=11,18) of 0.99 ± 0.05 and 1.08 ± 0.04 respectively. On CHOW diet, BMIs did not change after 3-month-, at 1.04 ± 0.05 (NTg, p=0.99 N=11) and 1.12 ± 0.04 (TgAD, p=1.00 N=18), or after 6-month-time frame, at 1.11 ± 0.10 (NTg, p=0.87 N=3) and 1.16 ± 0.04 (TgAD, p=0.88 N=4). In contrast, male NTg and TgAD rats fed with HCHF diet had a pronounced increase in BMI from 1.02 ± 0.02 (NTg) and 1.06 ± 0.04 (TgAD) at the onset of the diet regimen, to 1.25 ± 0.06 (p<0.01 N=14) and 1.27 ± 0.04 (p<0.01 N=18) after 3 months, then to 1.47 ± 0.04 (p<0.01 N=3) and 1.42 ± 0.07 (p<0.01 N=3) after 6 months of HCHF, respectively.

Female NTg and TgAD rats on CHOW had similar starting BMIs (p=0.41 N=13,11) of 0.53 ± 0.02 and 0.61 ± 0.02 respectively. Both female NTg (p<0.01 N=13,11) and TgAD (p<0.01 N=11,18) rats had much lower starting BMIs compared to males. Their BMIs did not change after 3-month-, at 0.59 ± 0.03 (NTg, p=0.92 N=13) and 0.65 ± 0.02 (TgAD, p=1.00 N=11), or after 6-month-time frame, at 0.63 ± 0.02 (NTg, p=0.92 N=4) and 0.76 ± 0.05 (TgAD, p=0.96 N=2). On the other hand, female NTg and TgAD rats markedly increased their BMI on HCHF diet, from BMI of 0.53 ± 0.02 and 0.60 ± 0.02, to 0.74 ± 0.03 (NTg, p<0.01 N=16) and 0.80 ± 0.04 (TgAD, p<0.01 N=17) after 3 months, then to 0.85 ± 0.07 (NTg, p<0.01 N=4) and 0.99 ± 0.06 (TgAD, p<0.01 N=3) after 6 months of HCHF diet. In summary, HCHF diet increased BMI of both male and female rats to the same extent; 3-month HCHF diet increased BMI of males by 0.21 and females by 0.20; 6-month HCHF increased BMI of males by 0.36 and females by 0.32.

### HCHF increases brain glucose uptake in female TgAD rats

One potential consequence of obesity induced by a varied HFHC diet is a systemically altered energy metabolism. Brain has a high demand for energy (Mergenthaler et al., 2013) thus changes in nutrient availability as a result of HCHF may alter overall metabolism and functioning of the brain. To evaluate metabolic changes, we used magnetic resonance spectroscopy (MRS) to measure uptake of 2-deoxy-D-glucose as a readout of glucose utilization within the hippocampus (HIPP), a region laden with AD pathology and crucial for cognitive function (Rao et al., 2022) (**Fig. 1A**). Three months of HCHF diet induced higher glucose uptake among rats of both genotypes (diet p=0.02), with no differences between genotypes (genotype p=0.33) (**Fig. 1B**) nor an interaction between genotype and diet (genotype*diet p=0.886) (**Fig. 1B**). Interestingly, examination of the sexes demonstrated that HCHF significantly increased glucose uptake in female TgAD but not male TgAD rats (sex*diet p=0.03) (**Fig. 1C**). These results suggest that obesity increased hippocampal glucose metabolism in female TgAD rats.

**Figure 1.**
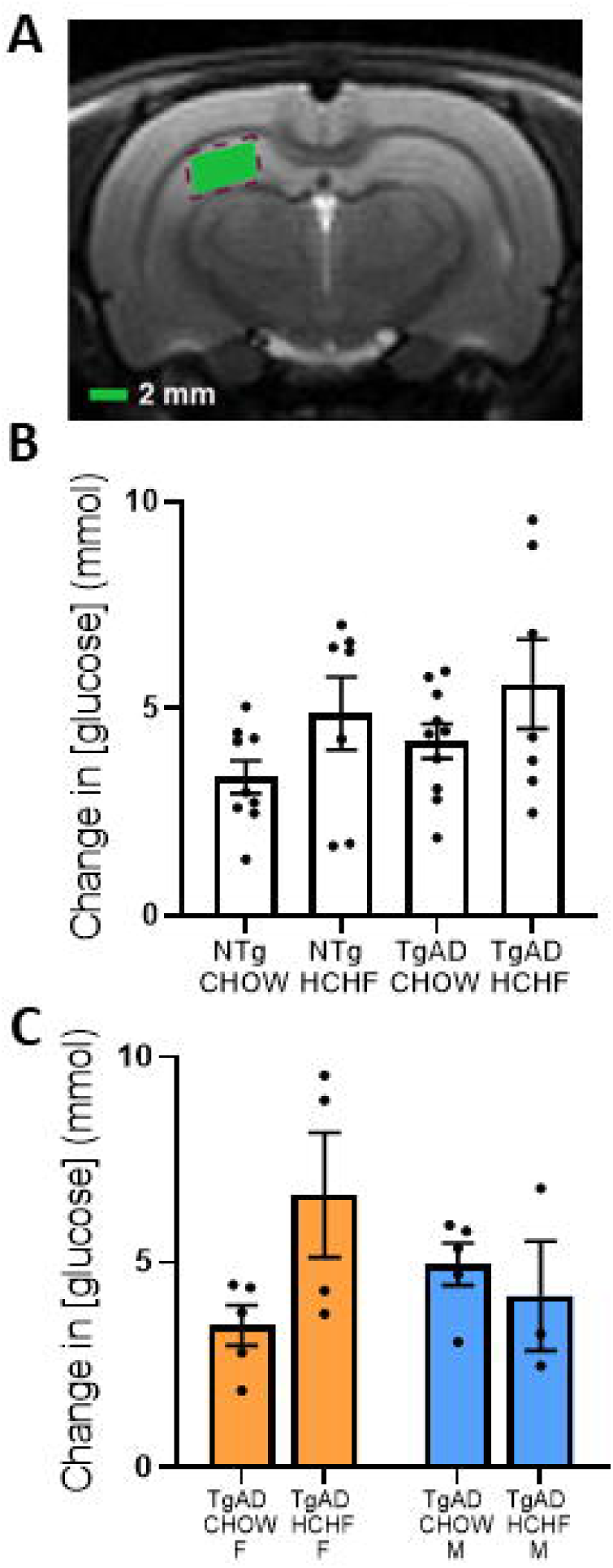
HCHF stimulates brain glucose metabolism in female TgAD rats. ***(A)*** T2-weighted coronal slice with shaded area representing the voxel used for MRS acquisition. ***(B)*** When both NTg and TgAD rats are considered, 3-month HCHF upregulated HIPP glucose uptake (diet p=0.02) without significant interaction with AD transgenes (genotype*diet p=0.89). Transgenes alone also did not affect glucose uptake (genotype p=0.33). ***(C)*** Sex-specific analysis of TgAD rats showed that HCHF stimulated glucose uptake in females but not males (sex*diet p=0.03). Sex alone and diet alone are also significant factors in altering glucose uptake (sex p=0.04; diet p<0.01). N = 4, 5, 5, and 4 for females and 5, 2, 5, and 3 for males respectively for NTg CHOW, NTg HCHF, TgAD CHOW, and TgAD HCHF.

### HCHF slows cognitive decline in TgF344-AD rats in a sex-dependant manner

After establishing that HCHF diet induced obesity and increased glucose metabolism, we investigated the effects of HCHF in comparison to CHOW fed NTg and TgAD rats on cognitive performance. We have previously shown cognitive deficits in TgAD rats at 12 months of age including deficits in spatial memory and executive function (Morrone et al., 2020). At the same age after consuming HCHF for 3 months, deficits in spatial memory (genotype p=0.02) and executive function (genotype p<0.01) attributed to AD transgenes were not affected by HCHF (genotype*diet p=0.25 and p=0.95 respectively) (**Fig. 2A-B**). In contrast, deficits in spatial learning (genotype p<0.01) were significantly improved (genotype*diet p<0.03) (**Fig. 2C**). In the absence of AD pathology, that is in NTg rats, HCHF did not affect spatial memory (p=0.81), executive function (p=0.12), or spatial learning (p=0.11) in NTg rats (**Fig. 2A-C**). When HCHF was extended to 6 months, interaction between AD transgenes and HCHF with respect to spatial learning was no longer significant (genotype*diet p=0.39), and there was no effect on spatial memory (genotype*diet p=0.91). Interestingly, with respect to executive function, after 6-months on HCHF, deficits caused by AD transgenes (genotype p=0.01) were ameliorated (genotype*diet p=0.03) (**Fig. 2D**). Overall, HCHF improved spatial learning at 3 months and executive function at 6 months.

**Figure 2.**
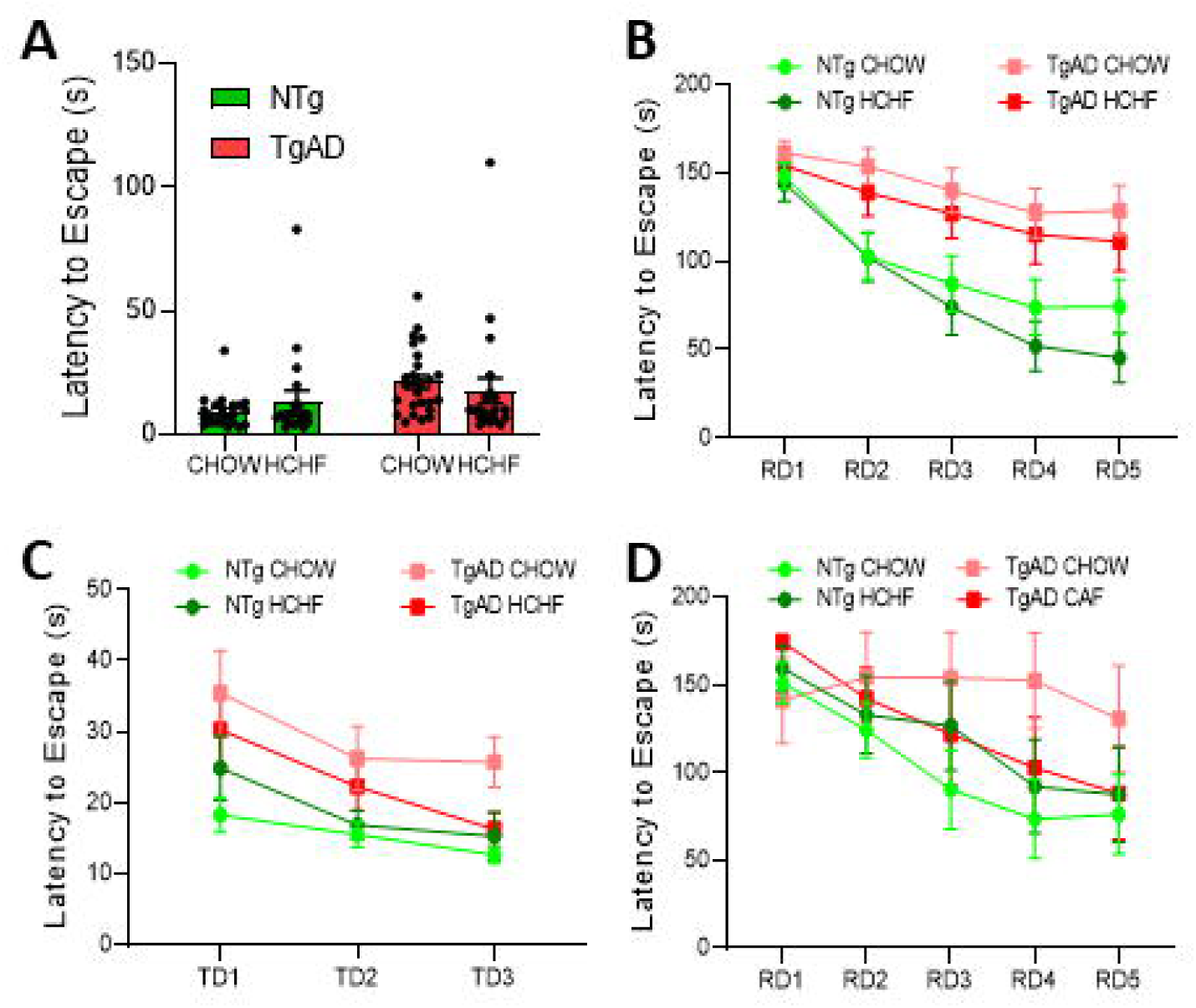
HCHF attenuates cognitive decline caused by AD transgenes. ***(A)*** At 12 months of age (3-month HCHF), impairment of spatial memory by AD transgenes (genotype p=0.02) was not attenuated by HCHF (genotype*diet p=0.25; diet p=0.92). ***(B)*** Deficits in executive function were induced by both AD transgenes and HCHF independently (genotype p<0.01; diet p=0.03) without significant interaction effects (genotype*diet p=0.95). ***(C***) Impairment of spatial learning by AD transgenes (genotype p<0.01) was slowed through interaction effects with HCHF (genotype*diet p=0.03; diet p=0.54). ***(D)*** At 15 months of age (6-month HCHF), deficit in executive function induced by AD transgenes (genotype p=0.01) was ameliorated by interaction with HCHF (genotype*diet p=0.03; diet p=0.86). For 3-month HCHF, N = 16, 12, 15, and 11 for females and 7, 8, 11, and 10 for males respectively for NTg CHOW, NTg HCHF, TgAD CHOW, and TgAD HCHF. For 6-month HCHF, N = 7, 7, 6, and 6 for the same experimental groups.

Since glucose uptake was increased in female but not in male TgAD rats, we hypothesized that their cognitive function might also be differentially affected. Thus, we then examined male and female TgAD rats separately and report remarkably different cognitive profiles in response to 3-month HCHF: With respect to spatial learning, male but not female TgAD rats trended towards improved performance in response to HCHF (sex*diet p=0.08) (**Fig. 3A**). While spatial memory was not affected by HCHF in either sex (sex*diet p=0.41), executive function was significantly improved by HCHF in female but not in male TgAD rats (sex*diet p=0.01) (**Fig. 3B**). Collectively, the results showed that changes to cognitive profiles of TgAD rats in response to HCHF are dependent on both time on the diet and sex.

**Figure 3.**
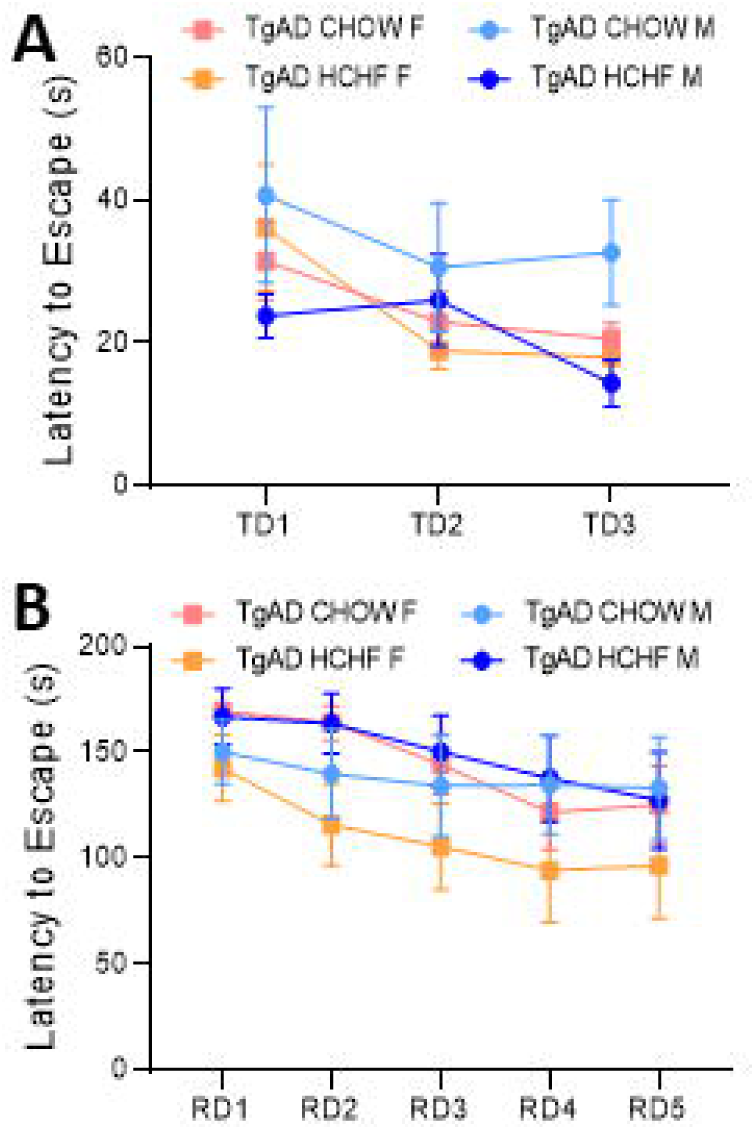
Rescue of spatial learning and executive function by HCHF is sex-specific. ***(A)*** Sex-specific analysis of TgAD rats showed that after 3-month HCHF, spatial learning in males but not females trended towards improvement (sex*diet p=0.08; sex p=0.36; diet p=0.06). ***(B)*** In contrast, executive function after 3-month HCHF improved in females but not males (sex*diet p=0.01; sex p=0.06; diet p=0.16). N = 16, 12, 15, and 11 for females and 7, 8, 11, and 10 for males respectively for NTg CHOW, NTg HCHF, TgAD CHOW, and TgAD HCHF.

### HCHF increases myelin density in female TgAD rats

We next examined whether the obesity-induced protection against cognitive decline correlated with structural changes at the tissue level. Myelin is a critical component of neuronal transmission and due to its high lipid content (Poitelon et al., 2020) and may directly be affected by HCHF-induced obesity. Preclinical studies have also shown that myelin density and integrity are tightly associated with AD progression: Myelin loss exacerbates AD pathology (Depp et al., 2023) while enhancing myelination improved cognitive deficits (Chen et al., 2021). We measured myelin density in prefrontal cortex (PFC) and hippocampus (HIPP) as these regions are important in spatial learning and executive function (Sigurdsson and Duvarci, 2015; Sampath et al., 2017).

In both PFC and HIPP, with respect to myelin density, 3-month HCHF had significant interaction effects (genotype*diet p<0.01 in PFC, p=0.01 in HIPP). Specifically, in NTg rats, HCHF decreased myelin (NTg CHOW vs. NTg HCHF p=0.04) to a level similar to that of TgAD CHOW rats (**Fig. 4A-B**) although this effect was not observed in HIPP (p=0.793) (**Fig. 4A-B**). In the absence of AD pathology, HCHF alone was thus detrimental to myelin in the PFC. TgAD CHOW rats had myelin loss compared to NTg CHOW rats (vs. TgAD CHOW p=0.06 in PFC, p<0.01 in HIPP) (**Fig. 4A-B**), but this myelin loss was mitigated by HCHF (TgAD CHOW vs. TgAD HCHF) in both PFC (p<0.01) and HIPP (p=0.01) (**Fig. 4A-B**). When HCHF was extended to 6 months, the interaction effect was no longer present (genotype*diet p=0.63 in PFC, p=0.07 in HIPP). Specifically, the effect of 6-month HCHF on NTg rats increased myelin density in PFC (NTg CHOW vs. NTg HCHF p=0.04) whereas the myelin rescue in TgAD rats was diminished (TgAD CHOW vs. TgAD HCHF p=0.16 for PFC, p=0.20 for HIPP) (**Fig. 4C**). We then looked at sex-specific outcomes and found that 3 months of HCHF increased myelin density in female (p<0.01 in PFC; p<0.01 in HIPP; sex*diet p=0.04 in PFC, 0.01 in HIPP) but not in male TgAD rats (p=0.56 in PFC, p=0.96 in HIPP) (**Fig. 4D**). These results are in line with our findings from executive function testing in that HCHF mitigates detrimental effects of AD transgenes specifically in females.

**Figure 4.**
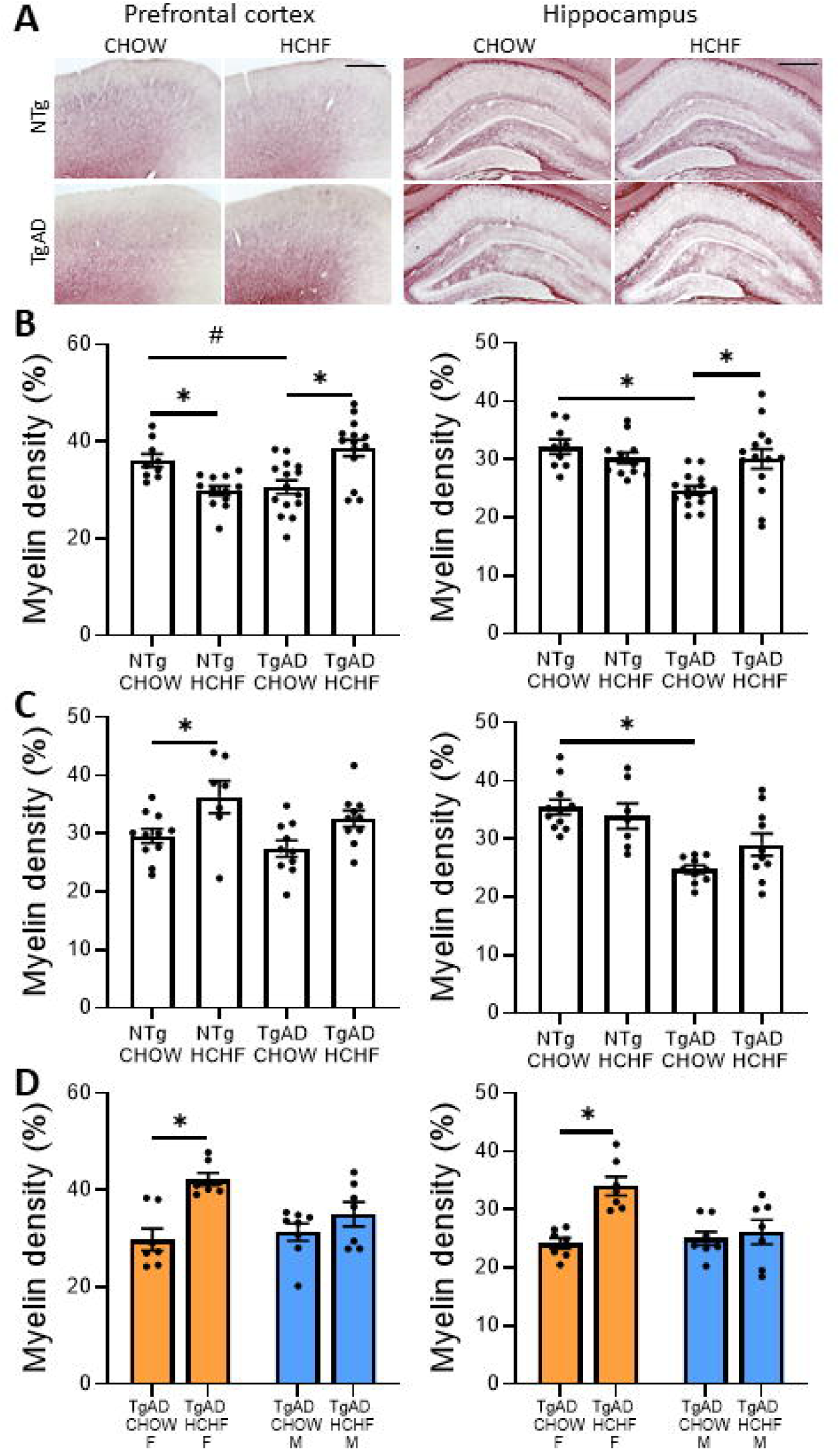
HCHF increases myelin density in female TgAD rats. ***(A)*** Representative images of Black Gold II-stained myelin in PFC (left, scale bar = 500 µm) and HIPP (right, scale bar = 800 µm). ***(B)*** HCHF for 3 months interacted with AD transgenes to increase myelin density of TgAD rats in both PFC (left, genotype*diet p<0.01; genotype p=0.27; diet p=0.55; TgAD CHOW vs. TgAD HCHF p<0.01) and HIPP (right, genotype*diet p=0.01; genotype p<0.01; diet p=0.17; TgAD CHOW vs. TgAD HCHF p=0.01). ***(C)*** HCHF for 6 months did not have interaction effects with AD transgenes with regards to myelin density in either PFC (left, genotype*diet p=0.63; genotype p=0.08; diet p<0.01; TgAD CHOW vs. TgAD HCHF p=0.12) or HIPP (right, genotype*diet p=0.07; genotype p<0.01; diet p=0.39; TgAD CHOW vs. TgAD HCHF p=0.20). ***(D)*** Sex-specific analysis of TgAD rats showed that 3-month HCHF increased myelin density in females but not males in both PFC (left, sex*diet p=0.04; sex p=0.17; diet p<0.01; CHOW F vs. CHOW HCHF p<0.01; CHOW M vs. CHOW HCHF p=0.56) and HIPP (right, sex*diet p=0.01; sex p=0.03; diet p<0.01; CHOW F vs. HCHF F p<0.01; CHOW M vs. HCHF M p=0.96). For 3-month HCHF, N = 5, 7, 7, and 7 for females and 4, 5, 8, and 7 for males respectively for NTg CHOW, NTg HCHF, TgAD CHOW, and TgAD HCHF. For 6-month HCHF, N = 11, 7, 10, 10 for the same experimental groups. * denotes p<0.05; # denotes p<0.10.

To further investigate myelin effects, we immunolabeled MBP which is a major component of CNS myelin that during disease and remodeling can either increase or decrease expression (Nam et al., 2018b; Lai et al., 2022). When both NTg and TgAD rats were considered, 3 months of HCHF increased MBP density in HIPP (diet p=0.57 for PFC, p<0.01 for HIPP) (**Fig. 5A**) but did not interact with genotype in either brain region (genotype*diet p=0.37 in PFC, p=0.33 in HIPP) (**Fig. 5A**). When HCHF was extended to 6 months, it continued to elevate MBP density in HIPP of both NTg and TgAD rats (diet p<0.01) (**Fig. 5B**). Furthermore, sex-specific effects of HCHF on MBP density resembled that of myelin density: In HIPP, 3-month HCHF resulted in MBP density trending towards an increase in female but not male TgAD rats (p=0.06 for females, p=0.79 for males; sex*diet p=0.22) (**Fig. 5C**). Overall, in agreement with the myelin increase, MBP density increased due to HCHF, indicating remodeling of myelin concurrent with myelin production.

**Figure 5.**
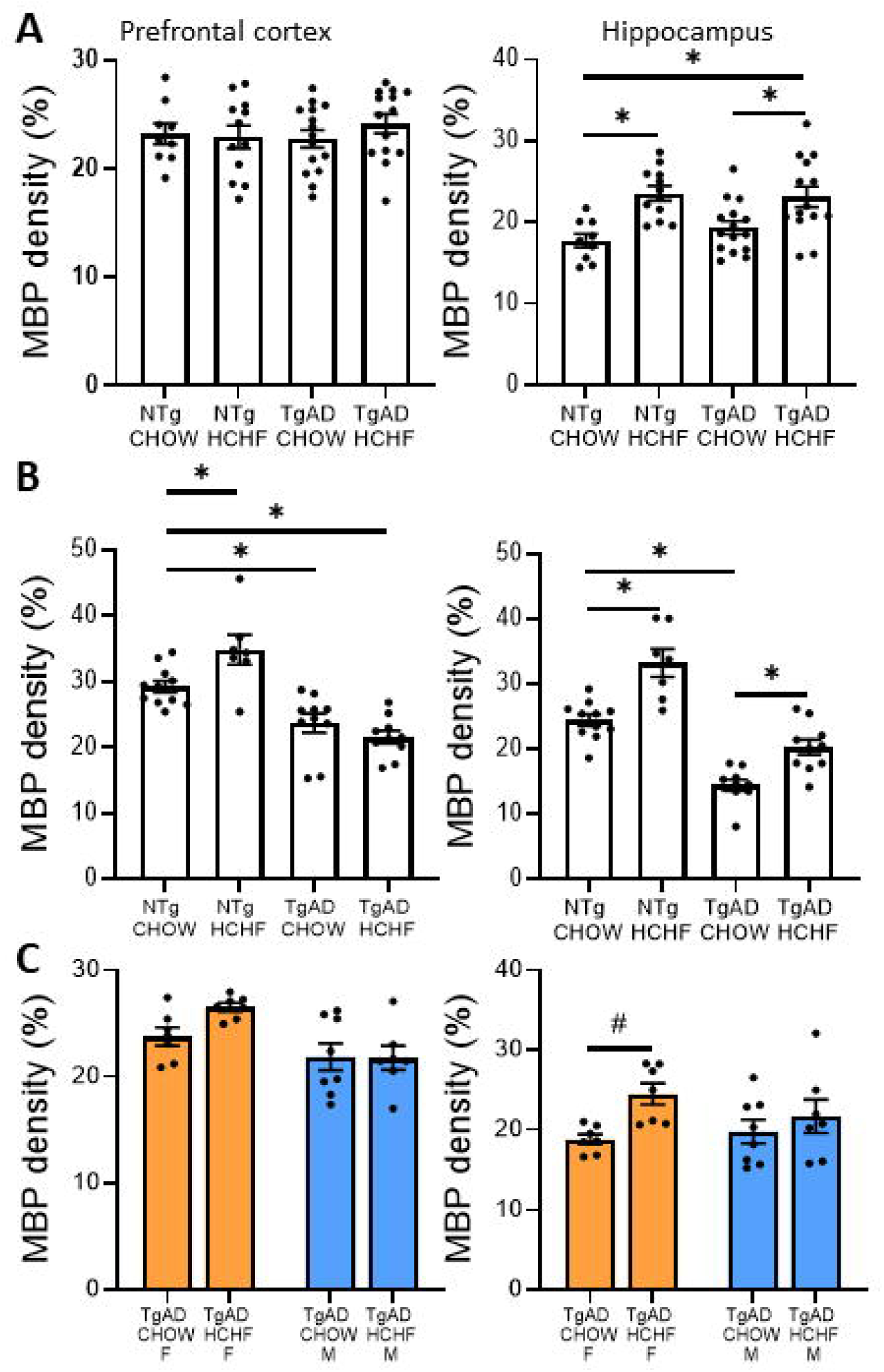
HCHF increases MBP density in both NTg and TgAD rats. ***(A)*** HCHF for 3 months did not affect MBP density in PFC (left, genotype*diet p=0.37; genotype p=0.69; diet p=0.57). In HIPP (right), HCHF increased MBP density of both NTg and TgAD rats (genotype*diet p=0.33; genotype p=0.58; diet p<0.01; NTg CHOW vs. NTg HCHF p<0.01; TgAD CHOW vs. TgAD HCHF p=0.03). ***(B)*** In PFC (left), 6-month HCHF increased MBP density in NTg but not TgAD rats (genotype*diet p=0.01; genotype p<0.01; diet p=0.21; NTg CHOW vs. NTg HCHF p=0.04; TgAD CHOW vs. TgAD HCHF p=0.66). In HIPP (right), 6-month HCHF increased MBP density in both NTg and TgAD rats (genotype*diet p=0.23; genotype p<0.01; diet p<0.01; NTg CHOW vs. NTg HCHF p<0.01; TgAD CHOW vs. TgAD HCHF p=0.01). ***(C)*** Sex-specific analysis of TgAD rats showed that 3-month HCHF did not have interaction effects with sex with regard to MBP density in either PFC (left, sex*diet p=0.16; sex p<0.01; diet p=0.20) or HIPP (right, sex*diet p=0.22; sex p=0.55; diet p=0.02). For 3-month HCHF, N = 5, 7, 7, and 7 for females and 4, 5, 8, and 7 for males respectively for NTg CHOW, NTg HCHF, TgAD CHOW, and TgAD HCHF. For 6-month HCHF, N = 11, 7, 10, 10 for the same experimental groups. * denotes p<0.05; # denotes p<0.10.

To investigate possible mechanisms by which HCHF diet might regulate myelin, we probed for oligodendrocytes, the myelin-producing cells in the brain, using the marker Olig2 and oligodendrocyte precursor cells (OPCs) to examine the number of myelinating cells, using the marker PDGFR-α (**Fig. 6A-B**). We found that with respect to oligodendrocyte density, 3-month HCHF diet resulted in significant interaction effects (genotype*diet p<0.01 in PFC, p=0.30 in HIPP) such that Olig2 density in TgAD HCHF rats compared to NTg CHOW rats was increased (p=0.047 in PFC, p<0.01 in HIPP) (**Fig. 6A, 7A**). In contrast, HCHF diet in NTg rats did not affect oligodendrocytes (NTg CHOW vs. NTg HCHF p=0.27 in PFC, p=0.93 in HIPP). When HCHF diet was extended to 6 months, interaction with AD transgenes was not significant (genotype*diet p=0.88 in PFC, p=0.67 in HIPP), and the beneficial effects on oligodendrocyte density in TgAD rats were lost (NTg CHOW vs. TgAD HCHF p=0.31 in PFC, p=0.59 in HIPP) (**Fig. 7B**). Interestingly, in TgAD rats fed for 3-months with HCHF, the increase in oligodendrocytes was detected only in males (p<0.01 in PFC, p=0.04 in HIPP) and not in females (p=1.00 in PFC, p=0.56 in HIPP; sex*diet p=0.02 in PFC, p=0.32 in HIPP) (**Fig. 7C**).

**Figure 6.**
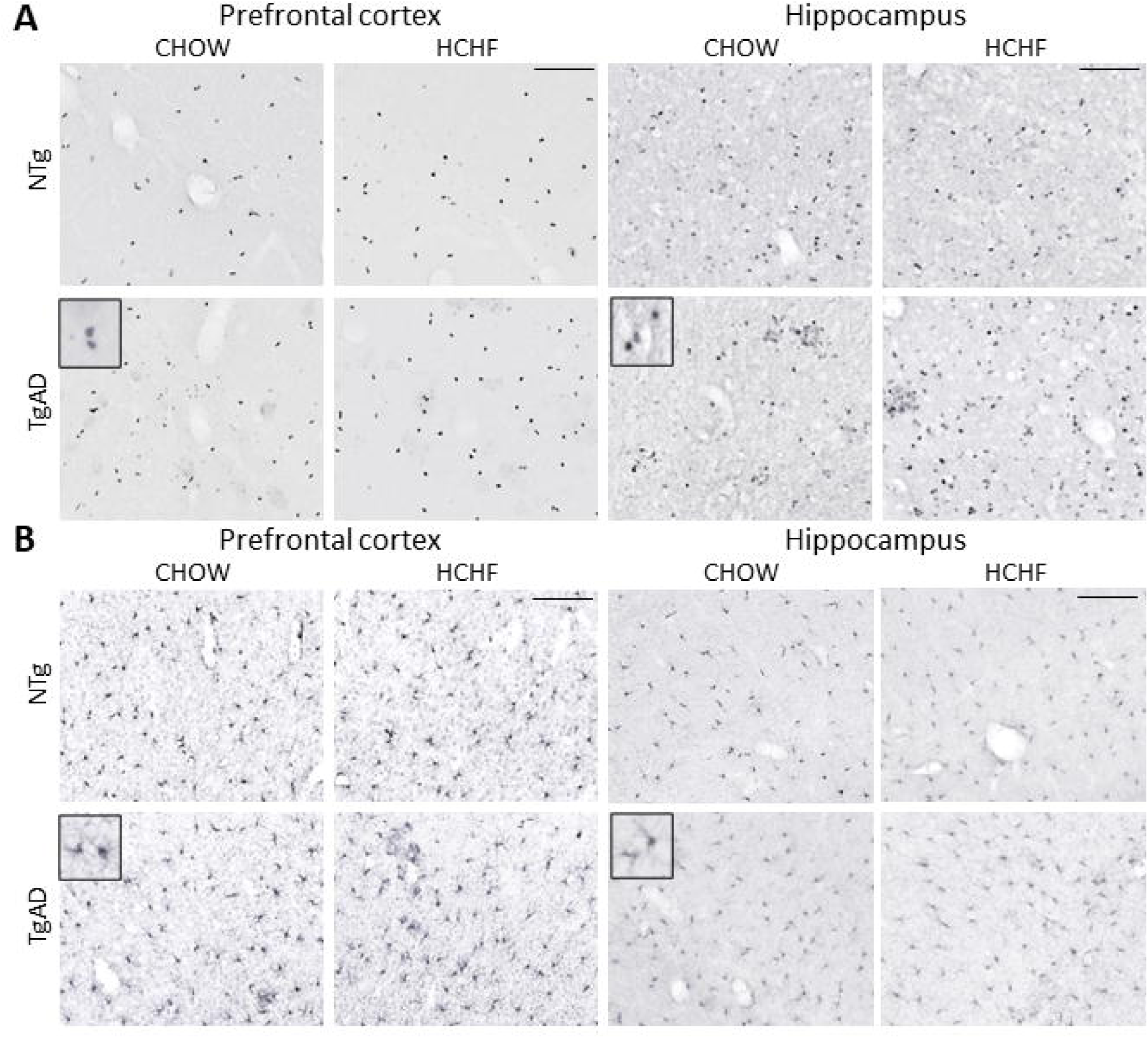
HCHF increases density of oligodendrocytes and OPCs in TgAD rats. ***(A)*** Representative images of Olig2-labeled oligodendrocytes in PFC (left, scale bar = 150 µm) and HIPP (right, scale bar = 150 µm). ***(B)*** Representative images of PDGFR-α-labeled OPCs in PFC (left, scale bar = 200 µm) and HIPP (right, scale bar = 200 µm).

**Figure 7.**
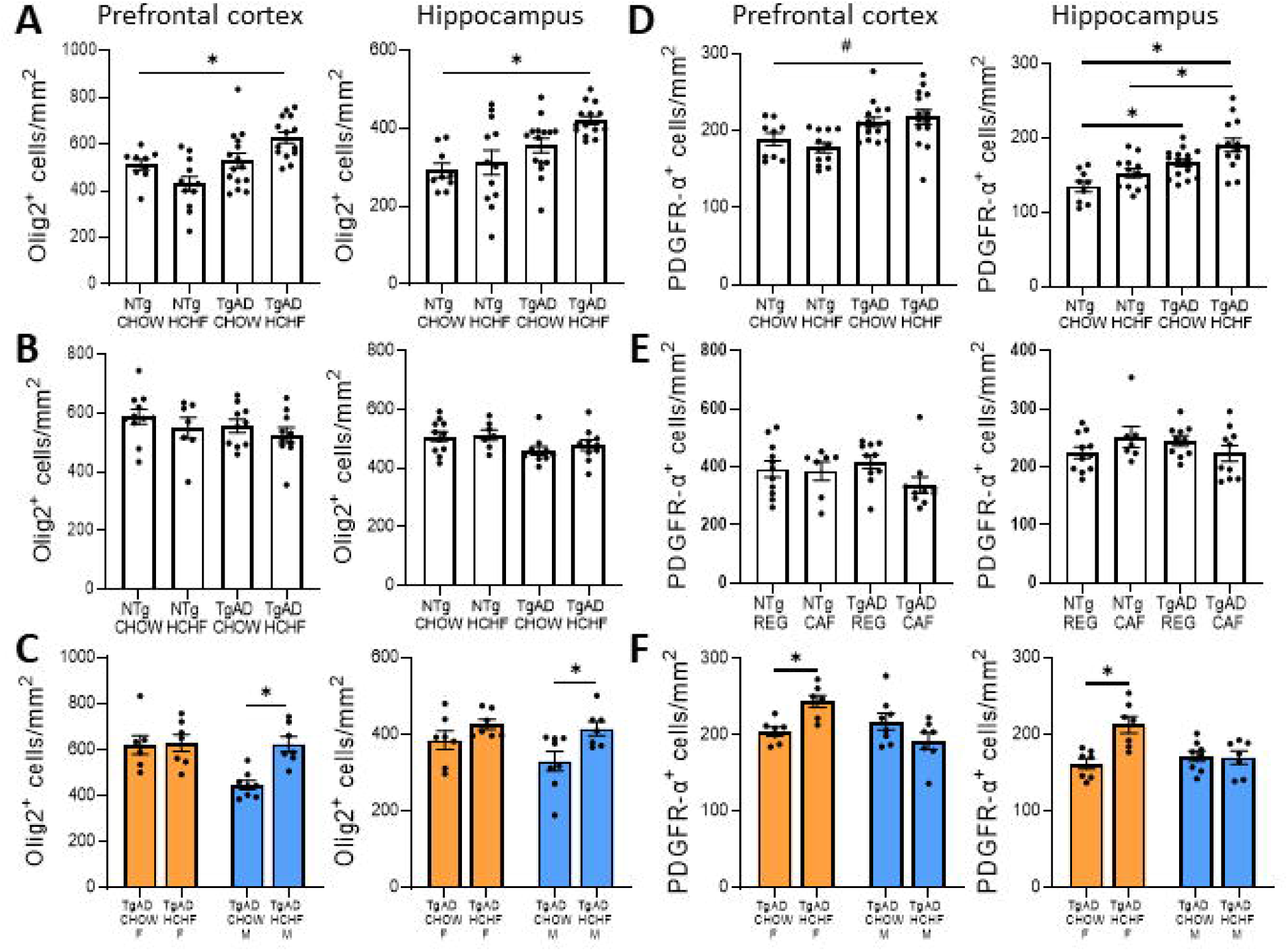
HCHF increases oligodendrocyte density in male TgAD rats and OPC density in female TgAD rats. ***(A)*** HCHF for 3 months increased oligodendrocyte density through interaction with AD transgenes in PFC (left, genotype*diet p<0.01; genotype p<0.01; diet p=0.77) but not in HIPP (right, genotype*diet p=0.30; genotype p<0.01; diet p=0.05). In both PFC (p=0.047) and HIPP (p<0.01), TgAD HCHF had significantly higher oligodendrocyte density compared to NTg CHOW. ***(B)*** HCHF for 6 months did not affect oligodendrocyte density in either PFC (left, genotype*diet p=0.88; genotype p=0.31; diet p=0.20) or HIPP (right, genotype*diet p=0.67; genotype p=0.02; diet p=0.48). TgAD HCHF rats no longer had higher oligodendrocyte density than NTg CHOW rats (p=0.31 in PFC, p=0.59 in HIPP. ***(C)*** Sex-specific analysis of TgAD rats showed that 3-month HCHF upregulated oligodendrocyte density in males and not females, in both PFC (left, sex*diet p=0.02; sex p=0.01; diet p=0.01; CHOW F vs. HCHF F p=1.00; CHOW M vs. HCHF M p<0.01) and in HIPP (right, sex*diet p=0.32; sex p=0.13; diet p=0.01; CHOW F vs. CHOW HCHF p=0.56; CHOW M vs. HCHF M p=0.04). ***(D)*** HCHF for 3 months increased OPC density in TgAD rats compared to NTg CHOW in both PFC (left, genotype*diet p=0.29; genotype p<0.01; diet p=0.79; NTg CHOW vs. TgAD HCHF p=0.09) and HIPP (right, genotype*diet p=0.65; genotype p<0.01; diet p=0.01; NTg CHOW vs. TgAD HCHF p<0.01). ***(E)*** HCHF for 6 months no longer increased OPC density in either PFC (left, genotype*diet p=0.20; genotype p=0.68; diet p=0.13; NTg CHOW vs. TgAD HCHF p=0.46) or HIPP (right, genotype*diet p=0.06; genotype p=0.76; diet p=0.83; NTg CHOW vs. TgAD HCHF p=1.00). ***(F)*** Sex-specific analysis of TgAD rats showed that 3-month HCHF upregulates OPC density in females and not males, in both PFC (left, sex*diet p<0.01; sex p=0.047; diet p=0.47; CHOW F vs. HCHF F p=0.03; CHOW M vs. HCHF M p=0.22) and HIPP (right, sex*diet p<0.01; sex p=0.07; diet p=0.01; CHOW F vs. HCHF F p<0.01; CHOW M vs. HCHF M p=1.00). For 3-month HCHF, N = 5, 7, 7, and 7 for females and 4, 5, 8, and 7 for males respectively for NTg CHOW, NTg HCHF, TgAD CHOW, and TgAD HCHF. For 6-month HCHF, N = 11, 7, 10, 10 for the same experimental groups. * denotes p<0.05; # denotes p<0.10.

We then examined OPCs using the marker PDGFR-α and found that although 3-month HCHF did not have interaction effects with AD transgenes (genotype*diet p=0.29 in PFC, p=0.65 in HIPP), HCHF in the presence of AD pathology increased OPC density (TgAD HCHF vs. NTg CHOW p=0.09 in PFC, p<0.01 in HIPP) (**Fig. 6A, 7D**), which mirrors the findings on oligodendrocytes. Similar to oligodendrocytes, lengthening HCHF to 6 months also diminished the effects on OPCs (genotype*diet p=0.20 in PFC, p=0.06 in HIPP; TgAD HCHF vs. NTg CHOW p=0.46 in PFC, p=1.00 in HIPP) (**Fig. 7D**). When examining sex differences, we found that in TgAD rats, 3-month HCHF increased number of OPCs in females (p<0.01 in PFC, p<0.01 in HIPP) but not in males (p=0.22 in PFC, p=1.00 in HIPP; sex*diet p<0.01 in PFC, p<0.01 in HIPP) (**Fig. 7F**) which is opposite to that observed with oligodendrocytes. Collectively, results showed that obesity in female TgAD rats prevented myelin loss due to AD progression and concurrently stimulated OPC proliferation, while obesity in male TgAD rats promoted oligodendrocyte maturation suggesting that obesity activated sex-dependent pathways in formation of oligodendrocytes.

### HCHF lowers NeuN in female TgAD rats

Increased myelin and oligodendrocytes or OPC densities may underlie improved cognitive performances in TgAD HCHF rats. As oligodendrocytes provide metabolic support for neurons (Philips and Rothstein, 2017), we then investigated whether neurons were preserved using the neuron-specific marker NeuN. Overall, 3-month HCHF diet did not yield interaction effects with AD transgenes (genotype*diet p=0.07 in PFC, p=0.40 in HIPP). Specific comparisons revealed, in comparison to CHOW fed rats, that HCHF did not alter NeuN density in either NTg (p=0.88 in PFC, p=0.78 in HIPP) or TgAD rats (p=0.21 in PFC, p=0.08 in HIPP); however, when comparing TgAD HCHF to NTg CHOW rats, a lowered NeuN density was surprisingly observed (p=0.03 in PFC, p<0.01 in HIPP) (**Fig. 8A-B**). Similar trends persisted after 6 months of HCHF diet: No interaction effects with AD transgenes were observed (genotype*diet p=0.80 in PFC, p=0.66 in HIPP). TgAD HCHF rats had lower NeuN density than that of NTg CHOW (p<0.01 in PFC, p=0.02 in HIPP). However, in both NTg and TgAD rats, 6-month HCHF decreased NeuN density to a greater extent (NTg CHOW vs. NTg HCHF p=0.01 in PFC, p=0.17 in HIPP; TgAD CHOW vs. TgAD HCHF p<0.01 in PFC; p=0.02 in HIPP) (**Fig. 8C**). In agreement with other outcome measures, response to HCHF was specific to female TgAD rats: In PFC, 3 months of HCHF lowered NeuN density in female rats (p=0.01) but not male rats (p=0.96; sex*diet p=0.01) (**Fig. 8D**); this sex-specific effect, however, was not significant in HIPP (sex*diet p=0.39) (**Fig. 8D**). Overall, these results show that in response to HCHF, NeuN density was lower in female TgAD rats, which may represent either neuronal dysfunction or death.

**Figure 8.**
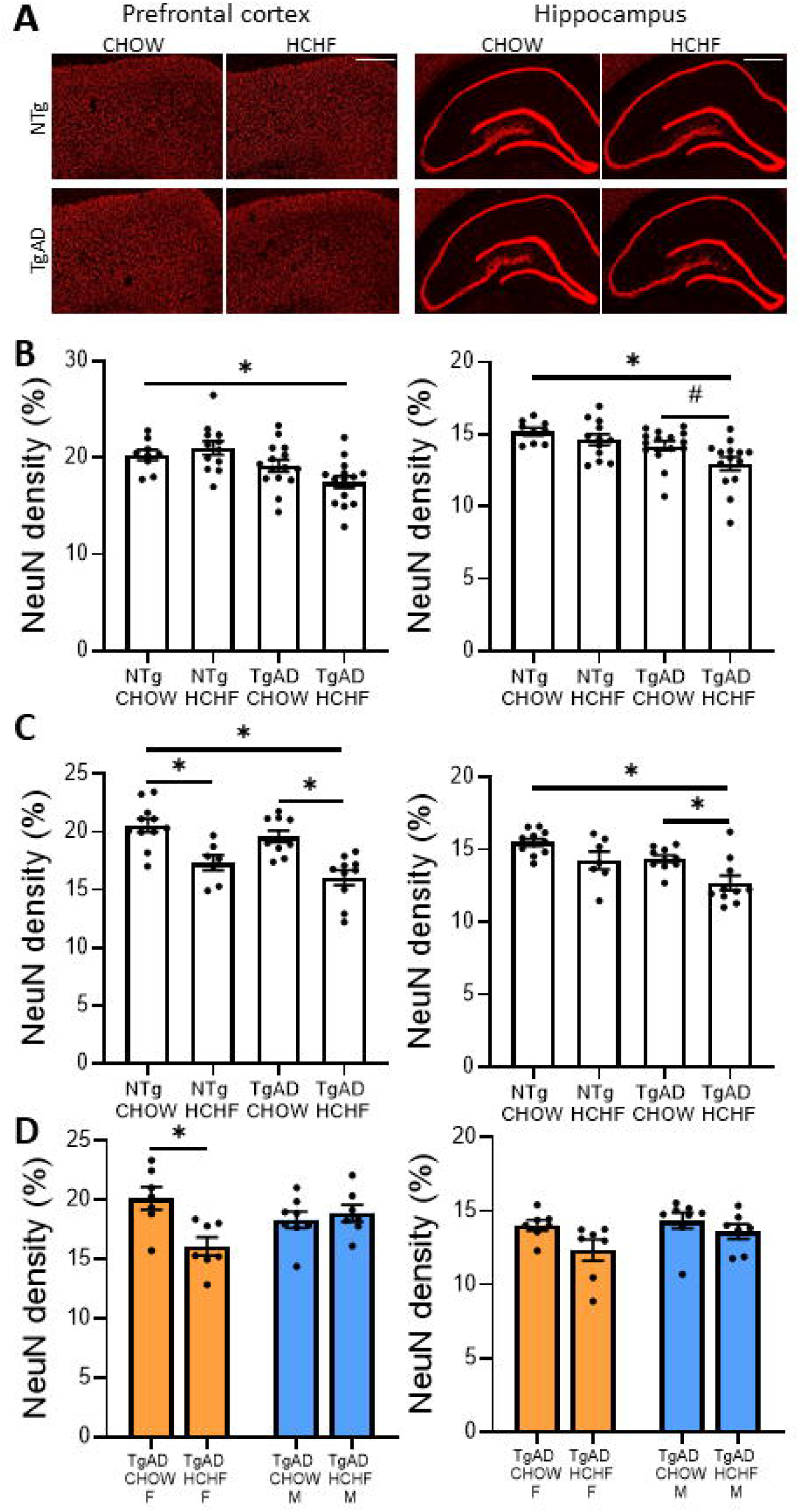
HCHF lowers NeuN density in female TgAD rats. ***(A)*** Representative images of NeuN-labeled neurons in PFC (left, scale bar = 500 µm) and HIPP (right, scale bar = 800 µm). ***(B)*** HCHF for 3 months decreased NeuN density in TgAD rats compared to NTg CHOW in both PFC (left, genotype*diet p=0.07; genotype p<0.01; diet p=0.48; NTg CHOW vs. TgAD HCHF p=0.03; TgAD CHOW vs. TgAD HCHF p=0.21) and HIPP (right, genotype*diet p=0.40; genotype p<0.01; diet p=0.03; NTg CHOW vs. TgAD HCHF p<0.01; TgAD CHOW vs. TgAD HCHF p=0.08). ***(C)*** HCHF for 6 months continued to lower NeuN density in TgAD rats compared to NTg CHOW in both PFC (left, genotype*diet p=0.80; genotype p=0.08; diet p<0.01; NTg CHOW vs. TgAD HCHF p<0.01; TgAD CHOW vs. TgAD HCHF p<0.01) and HIPP (right, genotype*diet p=0.61; genotype p<0.01; diet p<0.01; NTg CHOW vs. TgAD HCHF p<0.01; TgAD CHOW vs. TgAD HCHF p=0.02). ***(D)*** Sex-specific analysis of TgAD rats showed that in PFC (left), 3-month HCHF lowered NeuN density in females but not males (sex*diet p=0.01; sex p=0.54; diet p=0.04; CHOW F vs. HCHF F p=0.01; CHOW M vs. HCHF M p=0.96). This effect was not significant in HIPP (right; sex*diet p=0.39; sex p=0.16; diet p=0.04; CHOW F vs. HCHF F p=0.17; CHOW M vs. HCHF M p=0.77). For 3-month HCHF, N = 5, 7, 7, and 7 for females and 4, 5, 8, and 7 for males respectively for NTg CHOW, NTg HCHF, TgAD CHOW, and TgAD HCHF. For 6-month HCHF, N = 11, 7, 10, 10 for the same experimental groups. * denotes p<0.05; # denotes p<0.10.

### HCHF increases Aβ plaque load and tau inclusions

Lowered NeuN density in response to HCHF may have stemmed from altered accumulation of aggregated Aβ and tau, which are pathological hallmarks of AD present in our rat model (Morrone et al., 2022). Both Aβ and tau aggregates can cause neurotoxicity (Zhang et al., 2021); we thus examined whether HCHF effected levels of Aβ and tau aggregates. We measured Aβ plaques using 6F3D antibody, and after 3 months of HCHF or CHOW diet in TgAD rats, we found that HCHF diet significantly increased density of Aβ plaques in TgAD rats in PFC (p<0.01) but not in HIPP (p=0.79) (**Fig. 9A**). After 6 months of HCHF, this effect was diminished such that CHOW and HCHF TgAD rats did not have significantly different densities of Aβ plaques (p=0.58 in PFC, p=0.07 in HIPP) (**Fig. 9B**). Notably, when separated by sex, 3 months of HCHF increased Aβ plaque density only in PFC of female TgAD rats (p<0.01 in PFC, p=0.78 in HIPP; sex*diet p=0.01 in PFC; p=0.39 in HIPP) but in neither region in male TgAD rats (p=0.93 in PFC, p=0.96 in HIPP) (**Fig. 9C)**.

**Figure 9.**
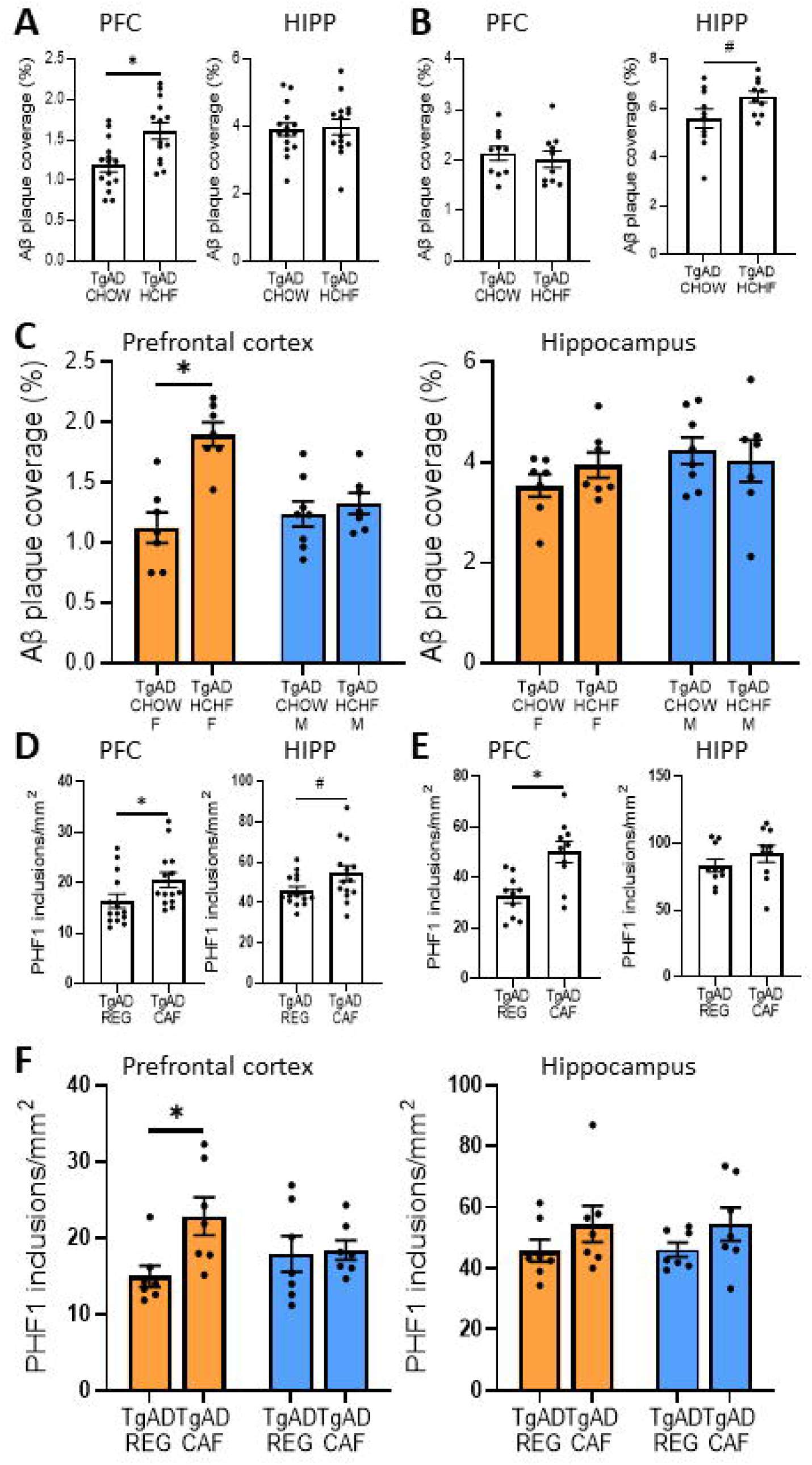
HCHF promotes aggregation of Aβ plaques and tau inclusions in female TgAD rats. ***(A)*** HCHF for 3 months increased density of Aβ plaques in PFC (left, p<0.01) but not in HIPP (right, p=0.79). ***(B)*** HCHF for 6 months no longer increased Aβ plaques in PFC (left, p=0.58). In HIPP (right), Aβ plaques trended towards increase in response to 6-month HCHF (right, p=0.07). ***(C)*** Sex-specific analysis showed that in PFC (left), 3-month HCHF increased density of Aβ plaques in females but not males (sex*diet p<0.01; sex p=0.04; diet p<0.01; CHOW F vs. HCHF F p<0.01; CHOW M vs. HCHF M p=0.93). This increase was not observed in HIPP (right, sex*diet p=0.32; sex p=0.20; diet p=0.73; CHOW F vs. HCHF F p=0.78; CHOW M vs. HCHF M p=0.96). For 3-month HCHF, N = 7 and 7 for females and 8 and 7 for males respectively for TgAD CHOW, and TgAD HCHF. For 6-month HCHF, N = 10 and 10 for the same experimental groups. ***(D)*** HCHF for 3 months increased density of PHF-1 tau inclusions in PFC (left, p= 0.049). In HIPP (right), tau inclusions trended towards increase after 3-month HCHF (p=0.07). ***(E)*** HCHF for 6 months increased density of PHF-1 tau in PFC (left, p<0.01) but not in HIPP (right, p=0.28). ***(F)*** Sex-specific analysis showed that in PFC (left), 3-month HCHF increased density of PHF-1 tau in females and not males (sex*diet p=0.07; sex p=0.70; diet p=0.04; CHOW F vs. HCHF F p=0.04; CHOW M vs. HCHF M p=0.998). In HIPP (right), HCHF did not affect tau density in either sex (sex*diet p=0.97; sex p=0.98; diet p=0.08; CHOW F vs. HCHF F p=0.55; CHOW M vs. HCHF M p=0.58). For 3-month HCHF, N = 7 for all experimental groups. For 6-month HCHF, N = 10. * denotes p<0.05; # denotes p<0.10.

To examine the role of tau pathology, we used PHF-1 antibody to label hyperphosphorylated tau inclusions in TgAD rats as a function of diet. After 3 months of HCHF, tau density increased in PFC (p=0.049) and trended towards increase in HIPP (p=0.07) (**Fig. 9D**). When HCHF was extended to 6 months, it continued to raise tau levels in PFC (p<0.01) but its effects in HIPP diminished (p=0.28) (**Fig. 9E**). When separated by sex, in PFC, 3-month HCHF diet increased tau in female (p=0.04) but not in male (p=1.00; sex*diet p=0.07) TgAD rats whereas in HIPP, neither sex had increased tau (p=0.55 in females, p=0.58 in males; sex*diet p=0.97) (**Fig. 9F**). These results collectively show that effects of HCHF on both Aβ plaques and tau inclusions sex-specifically target female TgAD rats, in parallel to their changes in myelin, OPCs, and neurons.

### HCHF dampens microglial activation

Aggregates of Aβ and tau promote inflammatory responses from microglia, the resident brain immune cells: They become activated as part of a mechanism to phagocytose and clear toxic protein species and cell debris, and such persisted activation is thought to be harmful to neurons (Leng and Edison, 2021). Microglia are also integral cells in maintenance of myelin through their regulation of oligodendrocyte maturation and pruning (McNamara et al., 2023). We thus investigated whether HCHF affected microglial density and activation using microglia-specific marker Iba1. After 3-month HCHF, we found significant interaction between diet and AD transgenes in PFC but not in HIPP (genotype*diet p=0.048 in PFC; p=0.34 in HIPP). Specifically, in the absence of AD pathology, HCHF did not alter microglial density (NTg CHOW vs. NTg HCHF p=1.00 in PFC, p=0.97 in HIPP) (**Fig. 10A-B**). TgAD CHOW rats have increased microglial density compared to NTg CHOW rats as a function of AD pathology (p=0.04 in PFC, p<0.01 in HIPP) (**Fig. 10A-B**). HCHF prevented this increase in TgAD rats in PFC (p=0.01) but not in HIPP (p=0.18) (**Fig. 10A-B**). Interestingly, HCHF decreased overall microglial density in TgAD rats to levels that were not significantly different from NTg CHOW in both PFC and HIPP (p=1.00 in PFC, p=0.07 in HIPP). When HCHF was extended for 6 months, microglial density was independent of interaction between genotype and diet (genotype*diet p=0.08 in PFC, p=0.81 in HIPP) in that CHOW and HCHF TgAD rats had similar densities of microglia (p=0.22 in PFC, p=0.93 in HIPP) (**Fig. 10C**). Furthermore, after 3 months of HCHF, sex and diet did not exert a significant interaction on microglial density in TgAD rats in either PFC (sex*diet p=0.68) or HIPP (sex*diet p=0.53) (**Fig. 10D**). Notably, HIPP microglial density was higher in males than in females regardless of diet (sex p=0.03) (**Fig. 10D**). It has been demonstrated that Iba-1 regulates actin-crosslinking involved in membrane ruffling, which is essential for the morphological changes that occur during microglia activation (Sasaki et al., 2001). Thus, based on our results, we propose that the decrease in Iba-1 density after 3 months of HCHF diet may be a result of decreased microglial activation.

**Figure 10.**
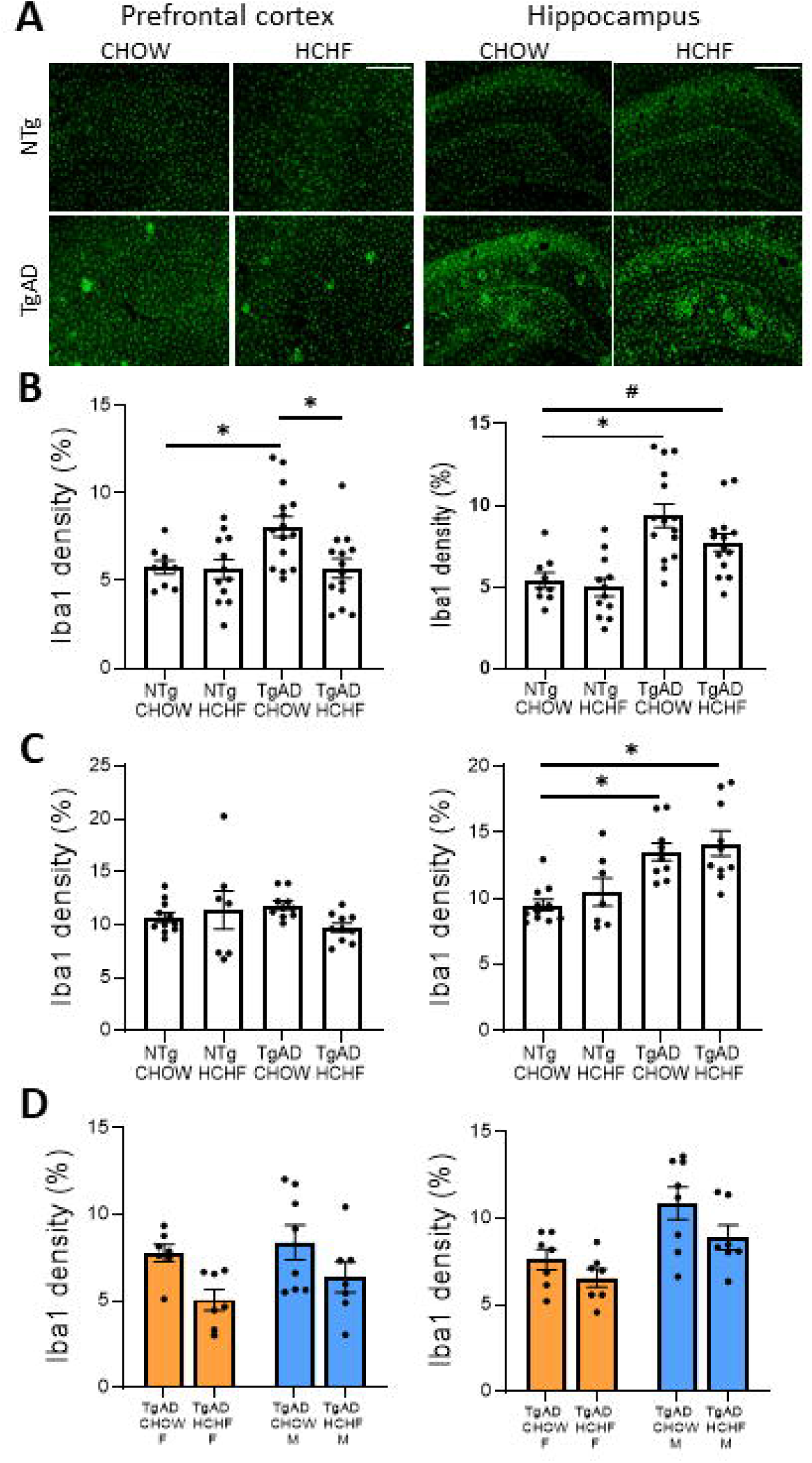
HCHF decreases microglial density in TgAD rats. ***(A)*** Representative images of Iba-1-labeled microglia in PFC (left, scale bar = 250 µm) and HIPP (right, scale bar = 500 µm). ***(B)*** In PFC (left), HCHF for 3 months prevents upregulation of microglial density by AD transgenes (genotype*diet p=0.048; genotype p=0.04; diet p=0.03) such that microglial density in TgAD HCHF rats was lower than that in TgAD CHOW (p=0.01) and was comparable to NTg CHOW (p=1.00). This effect was less pronounced in HIPP (right, genotype*diet p=0.34; genotype p<0.01; diet p=0.10; NTg CHOW vs. TgAD HCHF p=0.07; TgAD CHOW vs. TgAD HCHF p=0.18). ***(C)*** HCHF for 6 months no longer prevents increase of microglial density by AD transgenes in both PFC (left, genotype*diet p=0.08; genotype p=0.74; diet p=0.38; TgAD CHOW vs. TgAD HCHF p=0.22) and HIPP (right, genotype*diet p=0.81; genotype p<0.01; diet p=0.30; TgAD CHOW vs. TgAD HCHF p=0.93). ***(D)*** Separating males and females showed that 3-month HCHF did not affect microglial density in a sex-dependent manner in either PFC (left, sex*diet p=0.68; sex p=0.25; diet p=0.01) or HIPP (right, sex*diet p=0.53; sex p<0.01; diet p=0.047). For 3-month HCHF, N = 5, 7, 7, and 7 for females and 4, 5, 8, and 7 for males respectively for NTg CHOW, NTg HCHF, TgAD CHOW, and TgAD HCHF. For 6-month HCHF, N = 11, 7, 10, 10 for the same experimental groups. * denotes p<0.05; # denotes p<0.10.

It is now recognized that there are different populations of microglia depending on their proximity to disease/injury processes (Hawkes et al., 2012; Krasemann et al., 2017; Grubman et al., 2021). In light of this, we measured microglial activation in regards to plaque associated and non-associated populations through morphological analysis (**Fig. 11A**). After 3 months of HCHF, plaque-associated microglia had decreased level of activation in HIPP (p=0.01) but not in PFC (p=0.69) (**Fig. 11B**), whereas non-plaque-associated microglia were less activated in PFC (p<0.01) but not in HIPP (p=0.61) (**Fig. 11C**). Extending HCHF to 6 months did not alter plaque-associated microglia (p=0.12 in PFC, p=0.64 in HIPP) (**Fig. 11D**) but attenuated non-plaque-associated microglia in both regions (p<0.01 in PFC, p<0.01 in HIPP) (**Fig. 11E**). Further analyses on sub-populations showed no sex differences for plaque-associated microglia (sex*diet p=0.78 in PFC, p=0.85 in HIPP) and for non-plaque-associated microglia in HIPP (sex*diet p=0.66) in response to 3-month HCHF (**Fig. 11F**). In contrast, in PFC, HCHF significantly dampened activation of non-plaque-associated microglia in females (p=0.03) but not in males (p=0.33; sex*diet p=0.37) (**Fig. 11F**). This sex difference in PFC persisted when HCHF was extended to 6 months, where non-plaque-associated microglia was dampened in females (p<0.01) but not in males (p=0.25; sex*diet p=0.11) (**Fig. 11G**) whereas in HIPP, non-plaque-associated microglia were attenuated in both females (p=0.04) and males (p=0.04; sex*diet p=0.97). Altogether the results indicate that among the two microglial sub-populations, HCHF primarily dampens non-plaque-associated microglia particularly in female TgAD rats.

**Figure 11.**
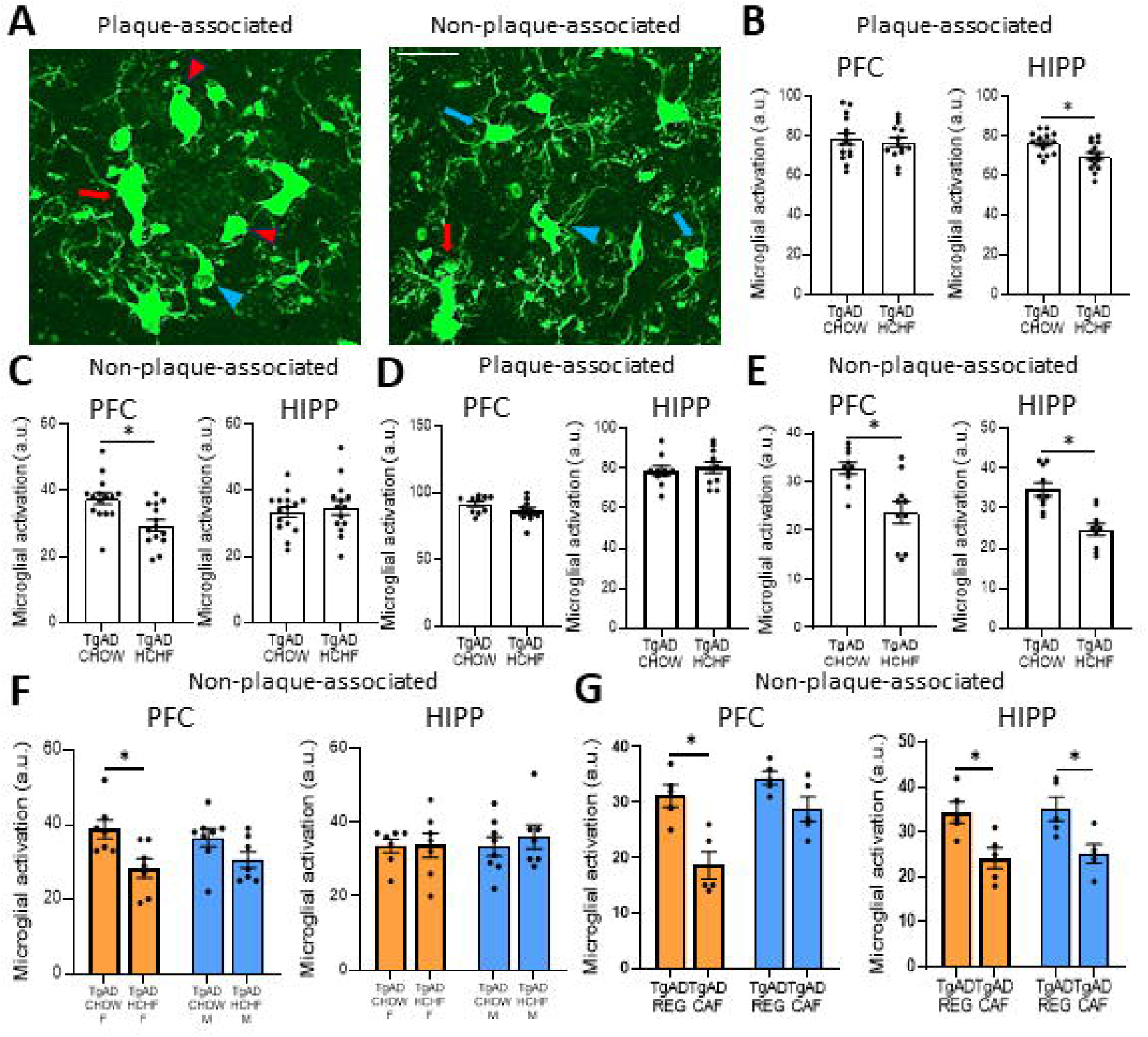
HCHF dampens microglial activation in female TgAD rats. ***(A)*** Representative images of Iba-1-labeled microglia that are either plaque-associated (left) or non-plaque-associated (right). Blue arrows, blue arrowheads, red arrows, and red arrowheads denote ramified, hypertrophic, bushy, and amoeboid morphologies respectively. Scale bar = 30 µm. ***(B)*** HCHF for 3 months dampened activation of plaque-associated microglia in HIPP (right, p=0.01) but not in PFC (left, p=0.69). ***(C)*** HCHF for 3 months also attenuated activation of non-plaque-associated microglia in PFC (left, p<0.01) but not in HIPP (right, p=0.61). ***(D)*** HCHF for 6 months did not affect activation of plaque-associated microglia in either PFC (left, p=0.12) or HIPP (right, p=0.64). ***(E)*** HCHF for 6 months dampened activation of non-plaque-associated microglia in both PFC (left, p<0.01) and HIPP (right, p<0.01). ***(F)*** In PFC (left), sex-specific analysis showed that 3-month HCHF attenuated activation of non-plaque-associated microglia in females but not males (sex*diet p=0.37; sex p=0.99; diet p<0.01; CHOW F vs. HCHF F p=0.03; CHOW M vs. HCHF M p=0.33). HCHF in HIPP (right) did not have the same effects (sex*diet p=0.66; sex p=0.71; diet p=0.63; CHOW F vs. HCHF F p=1.00; CHOW M vs. HCHF M p=0.90. ***(G)*** Sex-specific analysis of TgAD rats on 6-month HCHF also showed dampened activation of non-plaque-associated microglia in a female-specific manner in PFC (left, sex*diet p=0.11; sex p=0.01; diet p<0.01; CHOW F vs. HCHF F p<0.01; CHOW M vs. HCHF M p=0.25). In HIPP, dampening was observed in both sexes (right, sex*diet p=0.97; sex p=0.71; diet p<0.01; CHOW F vs. HCHF F p=0.04; CHOW M vs. HCHF M p=0.04). For 3-month HCHF, N = 7 and 7 for females and 8 and 7 for males respectively for TgAD CHOW, and TgAD HCHF. For 6-month HCHF, N = 10 and 10 for the same experimental groups. * denotes p<0.05.

## DISCUSSION

The current study aimed to further our understanding of the relationship between obesity and AD progression and elucidate potential mechanisms of their interaction. Collectively, our results show that obesity during the early stages of the disease protects against cognitive decline with sex-specific cognitive processes being affected. Females had improved executive function while males exhibited better spatial learning. Concurrent with ameliorated cognitive function in female TgAD rats are increased density of myelin and oligodendrocyte precursor cells and dampened activation of microglia suggesting that these changes likely contribute at least in part to improved cognition. In contrast, the underpinnings of the cognitive benefit of HCHF in males remain unclear. The cognitive benefits of obesity in females occur despite neuronal injury/loss and increased Aβ plaques and tau inclusions.

Here we report acceleration of Aβ plaque deposition and neuronal loss by obesity which agrees with previous studies using non-varied diets of high-fat or HCHF in rodent models of AD (Knight et al., 2014; Lin et al., 2016; Nam et al., 2017, 2018a; Sah et al., 2017; Walker et al., 2017; Medrano-Jiménez et al., 2019; Rollins et al., 2019; Bracko et al., 2020; Robison et al., 2020; Mazzei et al., 2021; Anderson et al., 2023). Importantly, we made novel observations of obesity concurrently driving neuroprotective processes in presence of accelerated AD pathology, including increased myelin and reduced activation of non-plaque-associated microglia. These seemingly dichotomous effects, all in females, have yet to be co-observed to date and provide important clues towards explaining the ‘obesity paradox’. Our dichotomous results show that timing of obesity with respect to AD progression and more importantly, sex, are key determinants in whether obesity exerts beneficial effects on disease progression. Notably, we have addressed two key issues with respect to timing of obesity: We have shown that the beneficial effects of obesity on myelin and microglia diminished with longer duration of disease; in contrast, neurotoxic processes including neuronal loss and increased tau aggregates persisted suggesting that moderating effects of obesity reach a threshold, whereby detrimental processes begin to outweigh the benefits. The other timing issue concerns whether obesity paradox is present during prodromal phase, early symptomatic phase, or full symptomatic phase of AD. Our results show that obesity during early symptomatic AD protects against cognitive decline.

It is somewhat surprising that obesity accelerated neuronal loss in females since their cognitive functions improved. Some neuronal loss may be attributed to increased aggregates of Aβ and tau. Nonetheless, our results demonstrated that maintaining density of myelin and OPCs in addition to increased glucose uptake in female TgAD rats was sufficient to overcome mild neuronal injury/loss such that cognitive dysfunction was prevented. Stimulating myelination through genetic or pharmacological means has been demonstrated to rescue cognitive decline in AD mice (Chen et al., 2021); similarly, obesity may have prevented cognitive loss through upregulation of myelin and OPCs. With respect to increased glucose uptake in females, it is probable that surviving neurons increased glucose metabolism as a compensatory response for mild neuronal loss. In clinical AD, changes in glucose metabolism correlate strongly with AD severity (Kumar et al., 2022). Increased glucose uptake observed here provide a feasible mechanism by which obesity lowers the risk of clinical AD. Another possibility is that synapses, whose integrity strongly correlates to cognitive maintenance (Jackson et al., 2019), remained intact due to dampening of activated microglia, which are known to excessively prune synapses in presence of AD pathology (Hong et al., 2016).

Higher load of Aβ and tau aggregates would be expected to stimulate proliferation and activation of microglia as the primary means of phagocytosing misfolded proteins in the brain (Nizami et al., 2019). We showed that the opposite happened in the presence of obesity particularly in non-plaque-associated microglia in female TgAD rats. Our results suggest that HCHF modulates deposition of Aβ and tau and non-plaque-associated microglia through independent pathways. Increased Aβ deposition with HCHF observed here is in agreement with results from previous studies that induced obesity in AD mouse models (Nam et al., 2017, 2018a; Medrano-Jiménez et al., 2019; Bracko et al., 2020; Mazzei et al., 2021). Studies of AD mice have also shown that obesity altered activity of Aβ-producing enzymes (Maesako et al., 2015; Bao et al., 2022), a probable mechanism by which Aβ deposition is increased. In our rat model, similar mechanisms may underlie observed increases in Aβ plaques and tau inclusions, but other independent processes are likely in play to concurrently dampen microglial activation.

Sex-specific effects of obesity on microglia observed in our study are particularly intriguing. We showed that obesity dampened activation of non-plaque-associated microglia only in females concurrent with female-specific improvement of executive function. Sex differences in AD have been understudied particularly in preclinical models of AD. As AD preferentially affects the female population, gaining mechanistic insights into sex differences is a priority. Obesity induces changes to the brain metabolic milieu, which underlies the sex-specific responses in female microglia as previous studies have demonstrated stark sex differences in metabolic responses of microglia in presence of AD: Microglia isolated from female AD mice upregulated several glucose metabolism enzymes towards a glycolytic phenotype (Guillot-Sestier et al., 2021). Post-mortem transcriptomic analysis of AD patients also showed drastic changes to metabolic pathways associated with cholesterol and steroids in female microglia (Hou et al., 2023). Although these findings show that female microglia have a metabolic profile more prone to dysregulation during AD, our results demonstrate that they are also more likely to benefit from metabolic changes induced by obesity.

The precise pathways by which obesity modifies metabolism in female microglia leading to neuroprotective phenotypes, and whether the same metabolic pathways in male microglia can be modulated accordingly, are key question to be addressed. Future therapeutic strategies can focus on reprogramming metabolic pathways harnessing the beneficial effects of obesity while avoiding its detrimental consequences. Metabolic reprogramming has been experimented both preclinically and in clinical trials: In streptozotocin-induced AD-like female rats, a panel of orally administered metabolic activators yielded beneficial effects on neurodegeneration (Turkez et al., 2023). The same metabolic activators also improved cognitive readouts in a Phase II clinical trial consisting of 69 AD patients (Yulug et al., 2023). Targeting metabolic pathways through reprogramming is thus a feasible approach for AD treatment.

In conclusion, we show that obesity induced through a varied HCHF diet stimulates multiple pathways in a time- and sex-dependent manner. Our results help to delineate the relationships between obesity and AD progression. We have identified non-plaque-associated microglia in females as a key player by which changes in metabolism can modulate function towards a beneficial disease-modifying phenotype. Identifying the specific metabolic pathways by which obesity changes each cell type in the brain will be the next important steps in laying the groundwork for metabolic reprogramming.

## ACKNOWLEDGEMENT

This work was funded by Canadian Institutes of Health Research Project Grant (grant number: PJY153101), Canada Research Chairs Program, National Institutes of Health R01 (grant number: AG057665-02), and Canadian Consortium on Neurodegeneration in Aging, which was supported by a grant from the Canadian Institute of Health Research with funding from several partners (grant number: CAN-137794), We thank Terrence Town and Tara M. Weitz for providing F344 rat breeding pairs.

